# Red Blood Cell Distribution Width: genetic evidence for aging pathways in 116,666 volunteers

**DOI:** 10.1101/128330

**Authors:** Luke C. Pilling, Janice L. Atkins, Michael O. Duff, Robin N. Beaumont, Samuel E. Jones, Jessica Tyrrell, Chia-Ling Kuo, Katherine S. Ruth, Marcus A. Tuke, Hanieh Yaghootkar, Andrew R. Wood, Anna Murray, Michael N. Weedon, Lorna W. Harries, George A. Kuchel, Luigi Ferrucci, Timothy M. Frayling, David Melzer

## Abstract

Variability in red blood cell volumes (distribution width, RDW) increases with age and is strongly predictive of mortality, incident coronary heart disease and cancer. We investigated inherited genetic variation associated with RDW in 116,666 UK Biobank human volunteers.

A large proportion RDW is explained by genetic variants (29%), especially in the older group (60+ year olds, 33.8%, <50 year olds, 28.4%). RDW was associated with 194 independent genetic signals; 71 are known for conditions including autoimmune disease, certain cancers, BMI, Alzheimer’s disease, longevity, age at menopause, bone density, myositis, Parkinson’s disease, and age-related macular degeneration. Pathways analysis showed enrichment for telomere maintenance, ribosomal RNA and apoptosis. The majority of RDW-associated signals were intronic (119 of 194), including SNP rs6602909 located in an intron of oncogene *GAS6*; the SNP is also an eQTL for this gene in whole blood. RDW-associated exonic genetic signals included a missense variant in *PNPLA3*, which codes for a triacylglycerol lipase, and a rare (1% frequency) deletion in *SMIM1*, involved in red blood cell formation.

Although increased RDW is predictive of cardiovascular outcomes, this was not explained by known CVD or related lipid genetic risks. The predictive value of RDW for a range of negative health outcomes may in part be due to variants influencing fundamental pathways of aging.

## Introduction

Increased variation in a person’s Red Blood Cell (RBC) volumes (RBC distribution width (RDW), also termed anisocytosis) is strongly predictive of a range of incident cardiovascular conditions, cancers and mortality [1]–[3]. Although RDW is routinely measured in clinical hematology reporting – it is calculated by dividing the standard deviation of mean cell volume (MCV) by the MCV and multiplying by 100, to yield a RDW percentage [4] – it is only used clinically for diagnosis of anemia subtypes. Understanding the mechanisms involved in the links between increased RDW and negative health outcomes could provide clues to potential interventions to improve prognosis in those with high RDW who are not anemic, particularly in older people.

Established clinical causes of increased RDW include anemia and other iron or folate deficiencies [5], dyslipidemia [6] and other metabolic abnormalities, and inflammation [7]. Proposed mechanisms for increased RDW also include impaired erythropoiesis (the generation of new RBC) perhaps due to effects of inflammation or senescence of erythropoietic cells in the bone marrow, plus variations in RBC survival [8]. A previous analysis of 36 blood cell traits identified genetic variants [9], but the genetic signals for RDW were not investigated in depth in relation to the mechanisms that might explain the predictive value of RDW for negative health outcomes in people with normal hemoglobin levels.

We aimed to investigate RDW (overall and excluding anemia) using genetic analysis in a large population cohort, to identify underlying mechanisms. This involved genome-wide analysis of associations to find independent signals, investigations of biological pathways implicated by the results, and overlap with known risk alleles. We also examined associations between RDW and known variant genetic risk score analysis for conditions predicted by RDW, including cardiovascular disease. For this analysis, we used the exceptionally large UK Biobank volunteer sample with standardized measures of RDW across the cohort.

## Results

We included 116,666 UK Biobank participants of white/British descent with complete hematology measures, covariate data, and genetics data from the interim data release (May 2015) in our analyses. The mean age was 57 years (SD: 7.9, min=40, max=70), with 52,541 aged 60 to 70 years old, and the majority (52.6%) were female: **Table 1**.

**Table 1.** Summary statistics for 116,666 UK Biobank participants

| **Trait** | **Mean (SD)** | **Min - Max** |
| --- | --- | --- |
| Age (years) | 56.92 (7.94) | 40 - 70 |
| RDW (%) | 13.49 (0.95) | 11.1 - 38.3 |
| Mean Cell Volume (fL) | 91.39 (4.45) | 54.5 - 160.3 |
| Hemoglobin conc. (g/dL) | 14.23 (1.23) | 0.14 - 20.5 |
| **Sex** | **N** | **%** |
| *Females* | 61,306 | 52.55 |
| *Males* | 55,361 | 47.45 |
| **Anemia** |  |  |
| *No* | 98,871 | 84.75 |
| *Yes \** | 17,795 | 15.25 |
| **RDW (%)** |  |  |
| *<12.5* | 8,703 | 7.46 |
| *12.5-12.9* | 24,559 | 21.05 |
| *13.0-13.4* | 33,804 | 28.97 |
| *13.5-13.9* | 25,413 | 21.78 |
| *14.0-14.4* | 12,967 | 11.11 |
| *14.5-14.9* | 5,460 | 4.68 |
| *≥15.0* | 5,761 | 4.94 |

\* = either hospital diagnosis or raised hemoglobin (WHO definition, see methods)

### Variance explained by genotypes

We estimated that 29.3% (SE = 0.5%) of the variance in RDW was accounted for by 457,643 directly genotyped variants with MAF>0.1%. In a secondary analysis we estimated the proportion explained in two groups: those aged ≥60 years (n=52,541), and those aged <50 years (n=24,988). The proportion of variance in RDW explained by the genetic variants was greater in the older group (33.8%, SE = 1.0%) compared to the younger group (28.4%, SE = 2.0%).

We estimated the proportion of variance in CHD (10,280 cases in 116,666 participants) accounted for by the variants to be 5.95% (SE = 0.45%). The proportion of the variance shared between RDW and CHD attributable to genetics is 6.62% (SE = 2.69%: 95% CIs = 1.35 to 11.9%).

### Genome-Wide Association Study

Of the 16,832,071 genetic variants included in this GWAS, 30,988 were significantly (p<5x10^-8^) associated with RDW (**Figure 1**; full results available to download here: http://www.t2diabetesgenes.org/data) after adjustment for age, sex, assessment center and array type (genetic relatedness is accounted for in the linear mixed models approach so no PCs are included – see methods). The 30,988 variants were mapped to 141 loci (runs of variants separated by <2Mb) on the genome, and included 194 independent signals after conditional analysis (“conditional SNPS”) (**Supplementary Table 1**).

**Figure 1.**
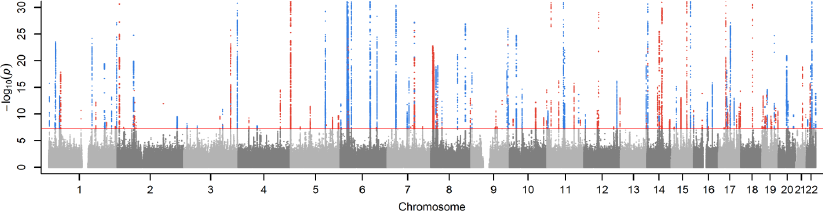
Genetic variants associated with RDW in GWAS of 116,666 UK Biobank participants. The variants are grouped into 194 independent signals, colored blue if a variant in the signal is associated with any trait in the NHGRI-EBI GWAS catalog of known associations, otherwise colored in red. The y-axis (–log_10_ *p*-values) is limited to 30 for clarity, as the max value is 200. See **Supplementary Table 1** for RDW associations for each signal, and **Supplementary Table 3** for mapping to the catalog. Horizontal line *p*=5x10^-8^.

After excluding 17,795 participants with anemia the GWAS of RDW was repeated on the 98,871 remaining participants; all 194 conditional SNPs remained nominally associated with RDW (p<0.001), but 24 were no longer genome-wide significant (p<5x10^-8^), possibly due to reduced power in the smaller sample size.

### Functional Implications of RDW-associated Genetic Variants

We utilized the UCSC Variant Annotation Integrator (http://www.genome.ucsc.edu/cgi-bin/hgVai) and the Ensembl Variant Effect Predictor (http://grch37.ensembl.org/Homo_sapiens/Tools/VEP) to interrogate a number of genomic annotation databases for the conditional SNPs. The majority (119 out of 194 total) were intronic, and 12 were located in the 3’ or 5’ un-translated regions (see **Supplementary Table 2** for complete variant annotation output). We found that 37 of 194 independent RDW-associated signals are known eQTLs (i.e. affect the expression of a gene) in whole blood (http://genenetwork.nl/bloodeqtlbrowser): 15 of these affect the gene predicted by the variant annotation integrator, including rs6602909, located in an intron of oncogene *GAS6* [10] (see **Supplementary Table 3** for complete matching of RDW-associated signals to eQTL data). SNP rs7775698 is an trans-eQTL for more than 30 genes.

**Table 2.** Four RDW-associated conditional genetic variants may have damaging effects on proteins

| Variant | Position |  |  | eA | Gene | AA | Codon | Effect | RDW association |  |  |
| --- | --- | --- | --- | --- | --- | --- | --- | --- | --- | --- | --- |
|  | CHR : POS | cDNA | Protein |  |  |  |  |  | Beta | P | eAF |
| SNP * |  |  |  |  |  |  |  |  |  |  |  |
| rs2075995 | 1:23847464 | 1105 | 226 | A | E2F2 | Q/H | caG/caT | P | 0.033 | 1.8x10<sup>-16</sup> | 0.502 |
| rs3811444 | 1:248039451 | 1169 | 374 | T | TRIM58 | T/M | aCg/aTg | D | 0.066 | 6.3x10<sup>-59</sup> | 0.333 |
| rs143845082 | 3:171417570 | 1539 | 398 | A | PLD1 | R/C | Cgt/Tgt | P | -0.217 | 4.6x10<sup>-13</sup> | 0.005 |
| rs149535568 | 3:171442535 | 1056 | 237 | A | PLD1 | G/C | Ggc/Tgc | D | -0.27 | 9.3x10<sup>-15</sup> | 0.048 |
| rs10479001 | 5:131607721 | 751 | 225 | T | PDLIM4 | A/V | gCa/gTa | B | -0.056 | 1.1x10<sup>-8</sup> | 0.042 |
| rs2578377 | 5:153413390 | 459 | 122 | T | FAM114A2 | G/S | Ggt/Agt | B | -0.026 | 4.1x10<sup>-10</sup> | 0.633 |
| rs1799945 | 6:26091179 | 347 | 63 | G | HFE | H/D | Cat/Gat | B | 0.138 | 2.2x10<sup>-143</sup> | 0.150 |
| rs368865 | 13:113479820 | 1037 | 317 | C | ATP11A | M/L | Atg/Ctg | B | -0.025 | 2.4x10<sup>-9</sup> | 0.724 |
| rs556052 | 19:49377436 | 1215 | 316 | C | PPP1R15A | A/P | Gct/Cct | B | 0.028 | 8.5x10<sup>-12</sup> | 0.333 |
| rs855791 | 22:37462936 | 2321 | 736 | T | TMPRSS6 | V/D | gTc/gAc | B | 0.119 | 1.2x10<sup>-204</sup> | 0.439 |
| rs738409 | 22:44324727 | 617 | 148 | G | PNPLA3 | I/M | atC/atG | D | 0.029 | 6.2x10<sup>-9</sup> | 0.216 |
| Deletion ¥ |  |  |  |  |  |  |  |  |  |  |  |
| rs566629828 | 1:3691997-3692014 | 309-325 | 21-26 | Del | SMIM1 | - | - | F | -0.12 | 4.5x10<sup>-12</sup> | 0.013 |

\* Output from UCSC Variant Annotation Integrator for the RDW-associated conditionally independent SNPs located in protein-coding regions. ¥ Output from Ensembl Variant Effect Predictor for insertion/deletion events.
“Effect”= (D)amaging, (P)ossibly damaing, or (B)enign SNP, or (F)rameshift due to deletion; “AA”=amino-acid change; “eA”=effect allele causing the change; “Beta”=the beta coefficient for the effect allele on RDW; “P”=p-value for the RDW association; “eAF”=effect allele frequency in UK Biobank white/British participants. All positions are from hg19/b37.

**Table 3.** MAGENTA results: biological pathways enriched in RDW genetics signals

| Biological pathway | N genes | Exp. Sig. | Obs. Sig. | p-value |
| --- | --- | --- | --- | --- |
| *Chromosomal & DNA-related* |  |  |  |  |
| Histone | 33 | 2 | 13 | 9.9x10<sup>-7</sup> |
| Nucleosome | 31 | 2 | 9 | 1.7x10<sup>-5</sup> |
| Nucleosome assembly | 43 | 2 | 10 | 1.0x10<sup>-4</sup> |
| Packaging of telomere ends | 23 | 1 | 11 | 9.9x10<sup>-7</sup> |
| Telomere maintenance | 49 | 2 | 12 | 5.0x10<sup>-6</sup> |
| Chromatin packaging and remodeling | 142 | 7 | 20 | 2.4x10<sup>-5</sup> |
| Apoptosis-induced DNA fragmentation | 9 | 0 | 4 | 4.0x10<sup>-4</sup> |
| *RNA polymerase-related* |  |  |  |  |
| RNA Pol I - promoter clearance | 44 | 2 | 14 | 9.9x10<sup>-7</sup> |
| RNA Pol I - promoter opening | 23 | 1 | 14 | 9.9x10<sup>-7</sup> |
| RNA Pol I - chain elongation | 30 | 2 | 9 | 7.0x10<sup>-6</sup> |
| RNA Pol I, III, and mitochondrial transcription | 82 | 4 | 15 | 1.2x10<sup>-5</sup> |
| *Lipid-related* |  |  |  |  |
| Chylomicron | 9 | 0 | 5 | 3.4x10<sup>-5</sup> |
| Chylomicron-mediated lipid transport | 15 | 1 | 5 | 6.0x10<sup>-4</sup> |
| Phospholipid efflux | 8 | 0 | 4 | 2.0x10<sup>-4</sup> |
| Very-low-density lipoprotein particle | 14 | 1 | 5 | 2.0x10<sup>-4</sup> |
| *Other* |  |  |  |  |
| Systemic Lupus Erythematosus | 60 | 3 | 15 | 9.9x10<sup>-7</sup> |
| Olfaction | 105 | 5 | 18 | 4.0x10<sup>-6</sup> |
| Cellular iron ion homeostasis | 25 | 1 | 8 | 2.8x10<sup>-5</sup> |
| Lactation (mammary development) | 8 | 0 | 3 | 4.5x10<sup>-3</sup> |

Output from MAGENTA GWAS enrichment software for the 30,988 RDW-associated genetic variants. Pathways shown are those where the estimated false discovery rate $p < 0.05$ . N genes=number of genes in the gene-set analysed; Exp. Sig.=Expected number of genes with a corrected gene p-value above the 95 percentile enrichment cutoff; Obs. Sig.=Observed number of genes with a corrected gene p-value above the 95 percentile enrichment cutoff; p-value=nominal p-value using 95 percentile of all gene scores for the enrichment cutoff.

Fifteen of the RDW-associated signals were exonic: 11 missense, 3 synonymous, and a 17 base-pair exonic deletion in gene *SMIM1*. The missense and deletion variants are shown in **Table 2**. PolyPhen-2 predicted that three missense variants are “probably damaging” to the protein function of genes *TRIM58, PLD1* and *PNPLA3*. Variant rs2075995 was predicted to be “possibly damaging” to the protein function of gene *E2F2*. *TRIM58* is a ubiquitin ligase induced during late erythropoiesis [10]; *PLD1* is a phospholipase implicated in processes including membrane trafficking [10]; *PNPLA3* is a triacylglycerol lipase in adipocytes and variant rs738409 is associated with susceptibility to Non-alcoholic Fatty Liver Disease [11]. Gene *SMIM1* is involved in RBC formation [10] and a 17 base-pair deletion (rs566629828) causing an exonic frameshift is strongly associated with increased RDW.

### GWAS Catalogue of Known Genetic Associations

Of the 194 conditional SNPs associated with RDW, 77 mapped to at least one trait in the catalogue of published GWAS (downloaded 13^th^ March 2017). This was arrived at by; filtering the 33,005 SNP-trait associations to those with p<5x10^-8^ (leaving 14,148 SNP-trait pairs for analysis); matching the positions to the UK Biobank results (13,146); filtering to those with significant RDW association in our analysis (p<5x10^-8^), leaving 923 SNP-trait pairs (420 unique SNPs; some SNPs are associated with multiple traits). These 420 unique SNPs mapped to 77 of the 194 conditional SNPs associated with RDW. These are shown in **Figure 1**; see **Supplementary Table 4** for further detail.

Traits also associated with the individual RDW variants included iron metabolism and several other red cell measures. Variants present were also associated with BMI, several lipids, hemoglobin A1C and metabolic syndrome, as well as height. Autoimmune associated conditions included autoimmune thyroid disease, type 1 diabetes, Crohn’s disease, inflammatory bowel disease, rheumatoid arthritis, systemic lupus erythematosus and ulcerative colitis. Variants linked to Lung, ovary and nasopharyngeal cancers were present. In addition, conditions associated with aging were represented, including variants linked to Alzheimer’s disease, age at menopause, bone density, Myositis, Parkinson’s disease, macular degeneration, C-reactive protein levels and longevity. For Alzheimer’s and longevity these were known SNPs in the *APOE* gene region.

### Gene Ontology Enrichment

MAGENTA software [12] identified pathways enriched in the genes mapped to variants significantly associated with RDW, including telomere maintenance, ribosomal RNA transcription and histone modifications (**Table 3**; **Supplementary Table 5**), plus apoptosis. In addition, pathways related to lipid metabolism (in particular chylomicrons) were also enriched.

### Genetic Risk Score associations

We tested 20 Genetic Risk Scores (GRS) for potentially explanatory traits for the predictive value of RDW for negative health outcomes; 7 were significant after adjustment for multiple testing (p<0.0025). Three were associated with raised RDW (HDL, type-1 diabetes, and BMI); four were associated with lower RDW (triglycerides, LDL, systolic blood pressure, and Alzheimer’s disease) (**Figure 2; Supplementary Table 6**). After exclusion of the ApoE locus the HDL GRS remained significant (p=0.003), but the AD GRS was no longer significant (p=0.84).

**Figure 2.**
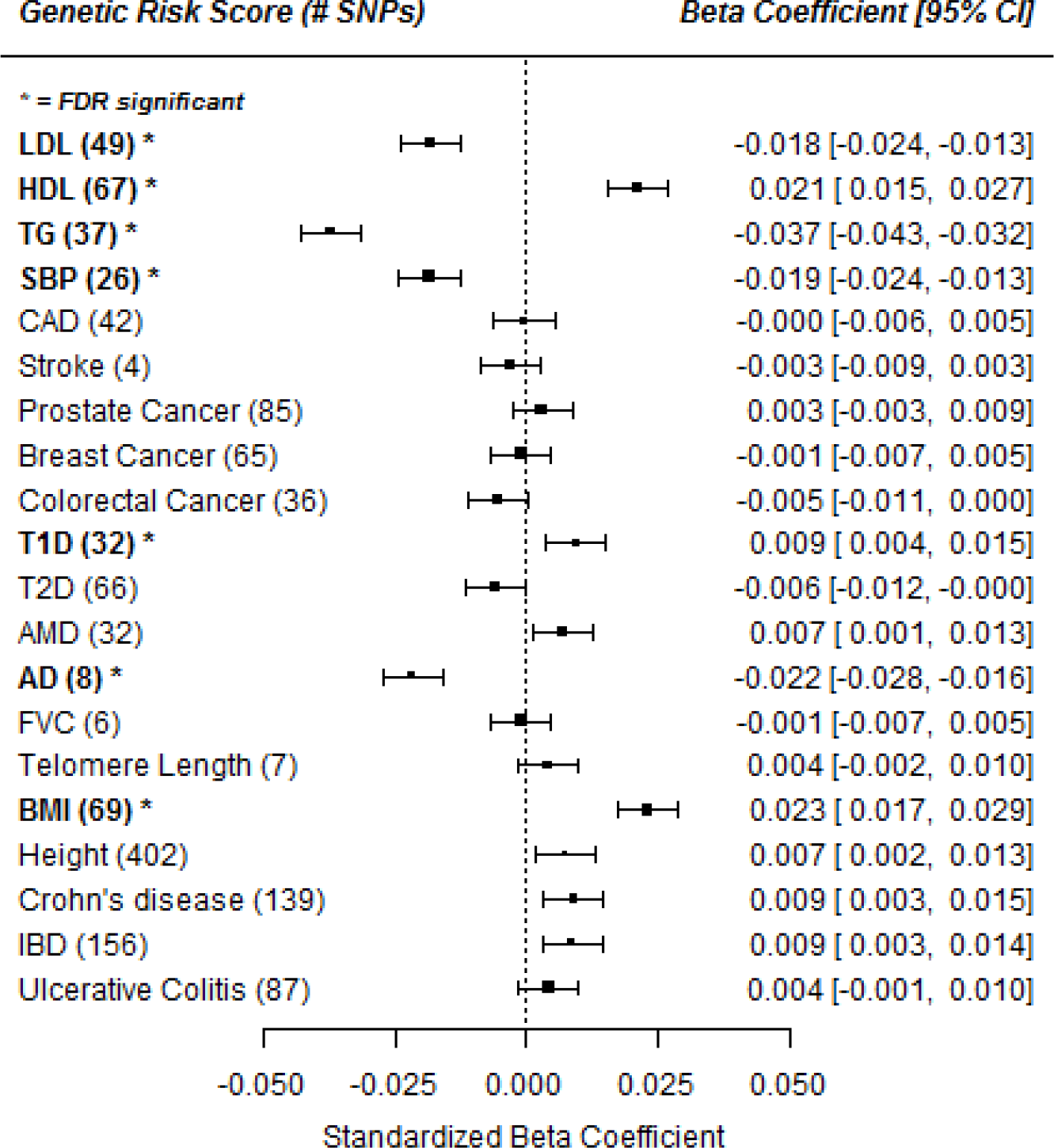
Genetic Risk Score associations with RDW. * FDR = false-discovery rate adjusted significant association. Genetic Risk Scores (GRS) were z-transformed prior to analysis. Linear regression model against RDW (z-transformed) including 116,666 participants, adjusted for age, sex, assessment center and population structure (genetic PCs 1-5). LDL (low-density lipoprotein), HDL (high-density lipoprotein), TG (triglycerides), SBP (systolic blood pressure), CAD (coronary artery disease), T1D (type-1 diabetes), T2D (type-2 diabetes), AMD (age-related macular degeneration), AD (Alzheimer’s disease), FVC (forced vital capacity), BMI (body mass index), IBD (inflammatory bowel disease). Full results in **Supplementary Table 5**.

Genetic risks for Crohn’s disease and inflammatory bowel disease were nominally associated with increased RDW (Beta=0.009: 95% CIs=0.003 to 0.015; Beta=0.009: 95% CIs=0.003 to 0.014; respectively). Risk scores for coronary artery disease were not associated with RDW, including after removal of the lipid related variants (Beta=0.00: 95% CIs=-0.006 to 0.005). A genetic risk score for telomere length was not significantly associated with RDW.

## Discussion

Variation in RBC size (RBC Distribution Width, RDW) increases markedly with age [13] and high RDW values are strongly predictive of increased mortality, plus incident cardiovascular disease and certain cancers [3], [14]. However, RDW is not generally considered as being a clinically useful measure outside the assessment of anemia sub-type, perhaps because the mechanisms explaining its prognostic value in people without anemia are unclear. In this study we investigated RDW using genetic analysis to understand the molecular mechanisms underpinning variation in RBC size.

A large proportion of RDW variation (29.3%) was attributable to common genetic variants in this analysis, and the variation explained by genetic variants appeared to increase with age, contrary to common assumptions that genetic effects decrease with advancing age. This could be due to effects accumulating over a time. Many of the “hallmarks of aging” also have this property [15]. Additionally, many of the RDW-associated genetic variants (in 71 of 194 conditionally independent signals) have previously been associated with other traits including metabolic syndrome, certain cancers, and autoimmune disease as well as aging related conditions including menopausal age.

Our analysis of genetic risk scores (GRS) showed that participants with genetically-increased risk for coronary artery disease or cancer did not have significantly higher RDW. Consistent with this, the proportion of the variance shared between RDW and coronary heart disease (CHD) attributable to genetics was only 6.6%; therefore the majority of genetically influenced red cell variation is independent of CHD.

As higher RDW is associated with CVD, we might have hypothesized that genetic risks for adverse lipid levels or blood pressures affect RDW, but instead we found associations in the opposite directions: GRS analysis showed that participants with genetically lower LDL levels, triglyceride levels, or systolic blood pressure, had higher RDW, and genetically higher HDL was associated with greater RDW. Published observational associations between lipids and RDW are only partially consistent with our findings here: Lippi *et al* found that LDL and HDL cholesterol were both negatively related to RDW, and triglyceride levels were positively related [6]. It is known that RBC have a role in cholesterol homeostasis by transporting cholesterol in the plasma membranes, with significant inter-individual differences not entirely explained by age or cholesterol levels [16]. The relationship between lipids and RDW is complex and requires further investigation. We also observed that genetically increased risk of type-1 diabetes was associated with increased RDW – further evidence for autoimmune involvement, in addition to the overlap in significant SNPs in the GWAS catalogue – and that genetically increased BMI is associated with increased RDW.

Genetic variants associated with RDW were enriched in expected pathways, including iron homeostasis, but we also found evidence for telomere maintenance, ribosomal RNA production, and a number of nucleosome and histone pathways. Short telomere length is a hallmark of cellular aging to senescence [15] and is associated with many risk factors of disease, however the causal direction is still uncertain [17], and longer telomeres have been linked to risk of cancer [18]. Kozlitina *et al* reported in 2012 that increased RDW is associated with shorter telomeres in leukocytes [19]. We created a genetic risk score for telomere length but it was not associated with RDW (Beta=0.004: 95% CIs=-0.002 to 0.010); further work is required to clarify this association.

We also found enrichment of RNA polymerase I (which transcribes ribosomal RNA) and RNA polymerase III (which transcribes transfer RNA); both are required for protein synthesis, including hemoglobin, and can even function as regulators of gene expression in their own right [20], suggesting these are key factors for consistent production of RBC. Deregulation of transcription and proteostasis are hallmarks of aging [15], and we have previously reported deregulation of gene expression of the transcriptional machinery with advancing age [21].

Four of the conditionally independent genetic variants associated with RDW are exonic and affect the amino-acid sequence of the protein products. Most others are intronic or intergenic, and may be regulatory; this is supported by published eQTL data [22], in which 37 of the RDW-associated signals have been reported to affect the expression of a gene in whole blood. These will be useful targets for future research to determine how these variants ultimately affect consistency in red blood cell size.

### Limitations

The UK Biobank is a volunteer study which achieved only a 5% response rate, so at assessment the participants were healthier than the general population. However, there was substantial variation in RDW within the participants so the results can still be generalized to the wider population [23].

More work is needed to establish the effects in other populations. No data have been released regarding the UK Biobank participant’s lipid and other blood assays; once this is available, further investigations into the complex relationship between RDW and lipids can be performed.

The GWAS catalogue does not contain every published GWAS, especially the most recent studies, but contains many of the largest meta-analyses for traits such as cardiovascular disease and cancer. It is therefore likely we have missed some studies using this method, therefore our results present an approximation of the overlap between RDW signals and other traits.

### Conclusions

Variation in RDW has a substantial genetic component, and this increases with increasing age. Although increased RDW is predictive of cardiovascular outcomes, this was not explained by known CVD or related lipid genetic risks. The predictive value of RDW for a range of negative health outcomes may in part be due to variants influencing fundamental pathways of aging.

## Methods

The UK Biobank study recruited 503,325 volunteers aged 40-70 who were seen between 2006 and 2010. Data includes RBC distribution width (RDW) and other clinical hematology measures, extensive questionnaires including smoking behavior and education history, and follow-up using electronic medical records. Currently one third of the participants have available genotype information. We utilized data from 116,666 participants of white/British descent with all available data.

### Phenotypes

RDW is a measure of the variability in the mean size of the RBC in each participant (in % units). It was measured using four Beckman Coulter LH750 instruments within 24 hours of blood draw, with extensive quality control performed by UK Biobank [24]. RDW is a continuous, highly skewed trait, therefore we used quantile normalization of the continuous measure to create a Gaussian distribution, so that the normality assumption of the linear regression models were not violated.

Anemia was determined both using self-reported diagnosis, electronic medical records (ICD10: D64* and D5* categories), or by low hemoglobin levels at the baseline assessment (<120g/L in females, <130g/L in males: from WHO definition [25]).

Coronary heart disease (CHD) was defined using self-reported myocardial infarction or angina, or diagnosis in the electronic medical records (ICD10: I20-I25).

### Variance explained and genetic correlation

We used BOLT-REML to determine the variance in RDW explained by the common, genotyped variants (n= 457,643 directly genotyped variants with MAF>0.1%, HWE p>1x10^-6^ and missingness <1.5%) using restricted maximum likelihood estimation [26]. BOLT-REML was also used to estimate the genetic correlation between RDW and CHD.

### Genome-wide association study

We performed a GWAS in 116,666 white/British participants (those with currently available genotyping from the 503,325 total UK Biobank participants), with complete genetic data to determine genetic variants associated with RDW. Over 800,000 genetic variants were directly genotyped using an Affymetrix Axiom array. After imputation, quality control, and filtering (we included autosomal variants with minor allele frequency (MAF) ≥0.1%, missingness <1.5%, imputation quality >0.4 and with Hardy-Weinberg equilibrium (HWE) p>1x10^-6^ within the white/British participants) 16,889,199 genetic variants were available for GWAS analysis: methods described in detail previously [27]–[29]. We also utilized data directly from the microarrays for variants on the X (n=19,381) and Y (n=284) chromosomes, and on the mitochondrial genome (n=135), which were unavailable in the imputed dataset.

GWAS was performed using BOLT-LMM, a software that uses linear mixed-effect models to determine associations between each variant and the outcome, incorporating genetic relatedness [26]. RDW residuals from a linear model adjusted for age, sex and assessment center were quantile-normalized prior to analysis. Models were adjusted for array type (two different Affymetrix arrays were used, which are >95% identical) at run time. Variants were classed as significant if the p-value for the association with RDW was less than 5x10^-8^.

### Identifying Conditionally Independent GWAS Signals

Many of the identified variants are correlated and may therefore not be independent; we used conditional analysis to determine independent signals by adjusting each variant in a locus for the most significant variant in that locus (loci defined as runs of SNPs separated by <2Mb on a chromosome). This process was repeated until only “conditional SNPs” remained that were significantly associated with RDW independent of one another.

Conditional SNPs were checked for their consistency of association with RDW in two sensitivity analyses: once excluding participants with prevalent anemia (either clinical diagnosis, self-report, or raised hemoglobin), and in a separate analysis excluding participants <60 years of age.

### GWAS-significant SNPs follow-up

The determine the gene-locations and possible functional consequences of the conditionally independent SNPs we submitted them to the UCSC Variant Annotation Integrator (https://genome.ucsc.edu/cgi-bin/hgVai), which combines information from several sources to determine the probable effect of a genetic variant, including on specific genes (for example intronic, missense, splice site, intergenic etc.) and PolyPhen-2 (a tool for predicting the impact of amino acid substitutions on the protein product). We used data from the “Blood eQTL browser” (http://genenetwork.nl/bloodeqtlbrowser) published by Westra *et al*. which reports associations between genetic variants and gene expression in whole blood [22] to determine whether the independent SNPs affect gene expression in human whole blood.

The GWAS catalogue of published variant-trait associations was searched for all SNPs (not just conditional SNPs) associated with RDW to determine which loci had previously been associated with another trait in GWAS analyses (p<5x10^-8^), and which were novel [30]. We used the UCSC `liftOver` tool (https://genome.ucsc.edu/cgi-bin/hgLiftOver) to match the genomic coordinates between the GWAS catalogue (GRCh38) and the UK Biobank genetics data (GRCh37).

We used the software package MAGENTA [12] to determine whether any biological pathways were enriched in the GWAS results, using the pathways database file GO_PANTHER_INGENUITY_KEGG_REACTOME_BIOCARTA.

### Genetic Risk Scores

Twenty Genetic Risk Scores (GRS) were computed for each participant based on the number of trait-raising alleles they have for a particular phenotype, such as LDL cholesterol or type-2 diabetes. These were computed according to the method described in Pilling *et al* 2016 [29]. Each of the 20 GRS computed was tested for its association with RDW using linear regression models, adjusted for age, sex, assessment center, genotype array, and population stratification (using the first 5 principal components (PCs)).

## Funding and Acknowledgements

This work was supported by an award to DM, TF, AM, LH and CB by the UK Medical Research Council (grant number MR/M023095/1). SEJ is funded by the Medical Research Council (grant: MR/M005070/1). JT is funded by a Diabetes Research and Wellness Foundation Fellowship. R.B. is funded by the Wellcome Trust and Royal Society grant: 104150/Z/14/Z. MAT, MNW and AM are supported by the Wellcome Trust Institutional Strategic Support Award (WT097835MF). RMF is a Sir Henry Dale Fellow (Wellcome Trust and Royal Society grant: 104150/Z/14/Z). ARW, HY, and TMF are supported by the European Research Council grant: 323195:GLUCOSEGENES-FP7-IDEAS-ERC. The funders had no influence on study design, data collection and analysis, decision to publish, or preparation of the manuscript. LF is supported by the Intramural Research Program of the National Institute on Aging, U.S. National Institutes of Health. Input from MD, CL and GK was supported by the University of Connecticut Health Center.

This research has been conducted using the UK Biobank Resource under Application Number 14631. We thank UK Biobank participants and coordinators for this dataset.

